# Nonself Mutations in the Spike Protein Suggest an Increase in the Antigenicity and a Decrease in the Virulence of the Omicron Variant of SARS-CoV-2

**DOI:** 10.1101/2021.12.30.474613

**Authors:** Joji M. Otaki, Wataru Nakasone, Morikazu Nakamura

## Abstract

Despite extensive worldwide vaccination, the current COVID-19 pandemic caused by SARS-CoV-2 continues. The Omicron variant is a recently emerged variant of concern and is now taking over the Delta variant. To characterize the potential antigenicity of the Omicron variant, we examined the distributions of SARS-CoV-2 nonself mutations (in reference to the human proteome) as 5 amino acid stretches of short constituent sequences (SCSs) in the Omicron and Delta proteomes. The number of nonself SCSs did not differ much throughout the Omicron, Delta, and Reference Sequence (RefSeq) proteomes but markedly increased in the receptor binding domain (RBD) of the Omicron spike protein compared to those of the Delta and RefSeq proteins. In contrast, the number of nonself SCSs decreased in non-RBD regions in the Omicron spike protein, compensating for the increase in the RBD. Several nonself SCSs were tandemly present in the RBD of the Omicron spike protein, likely as a result of selection for higher binding affinity to the ACE2 receptor (and hence higher infectivity and transmissibility) at the expense of increased antigenicity. Taken together, the present results suggest that the Omicron variant has evolved to have higher antigenicity and less virulence in humans despite increased infectivity and transmissibility.

## 1. Introduction

Despite worldwide efforts for vaccination, the COVID-19 pandemic still prevails as of December 2021, two years after its pathogenic emergence caused by a novel coronavirus, SARS-CoV-2 [1–4]. Recently, a new variant emerged from South Africa, which was announced on 25 November 2021 [5] and was designated the Omicron variant (B.1.1.529), one of the variants of concern announced by the World Health Organization (WHO) on 26 November 2021 [6]. A risk assessment of the Omicron variant was urgently released on 2 December 2021 [7]. Currently, the Omicron variant is spreading worldwide, including in Denmark [8] and the United States of America [9], displacing a previous variant of concern, the Delta variant (B.1.617.2) [6]. The Omicron variant contains a large number of unique mutations and appears to have higher infectivity and transmissibility than previous variants, raising a public health concern [10–13]. Prompt characterization of the Omicron variant is of high importance for public health.

Possible functional changes associated with mutations in the Omicron variant have already been evaluated by several computational studies. Several mutations have been localized in the receptor binding domain (RBD) of the spike (S) protein, possibly contributing to higher affinity to the ACE2 receptor and lower affinity to pre-existing infection-induced or vaccine-induced antibodies [13–15]. An increase in hydrophobic amino acid residues [15] or in electrostatic interactions [16,17] introduced by mutations has been linked to potentially higher infectivity and transmissibility. Computational analyses, including a structural analysis [13], an analysis based on artificial intelligence (AI) trained with numerous experimental data [18], and an analysis with amino acid interaction (AAI) networks [19], have revealed potential resistance of the Omicron variant against pre-existing antibodies. A large-scale SARS-CoV-2 genome analysis has suggested that Omicron mutations have been selected for vaccine resistance [20]. Consistent with these in silico studies, in vitro functional analyses of the Omicron variant have suggested significant escape from pre-existing infection-induced or vaccine-induced antibodies [21,22]. However, pre-existing T-cell immunity has still been effective [22,23]. These studies have already addressed some of the concerns associated with the Omicron variant, but alternative and complementary methods to further characterize new variants will also be helpful. Here, we employed a novel method simply based on amino acid sequences of SARS-CoV-2 and human proteomes to evaluate the antigenicity of the Omicron variant.

The human immune system recognizes foreign proteins based on short amino acid sequences presented as peptides by MHC molecules [24–26]. In the present study, a stretch of an amino acid sequence, when it exists as a part of a protein, is called a short constituent sequence (SCS; pronounced as [es/si:/es] or [ʃᵓks|). Operationally speaking, the human immune system stores memories of all possible SCSs from the human proteome, which are here called self SCSs. Every peptide presented by MHC molecules is collated with a dataset of self SCSs. When a sequence of a given peptide presented by MHC molecules is found in the dataset, it is recognized as “self”, a part of a human body. In contrast, when a sequence of a peptide is not found in the dataset, it is recognized as “nonself”, a foreign object to be eliminated by the immune system. These SCSs in a protein are called nonself SCSs, and they are antigenic by definition, although their antigenicity in vitro and in vivo should be determined experimentally. When a given peptide sequence increases rapidly to emergency levels when a microbe or virus infects a human body, the immune system likely produces antibodies against it even if it is found in a dataset. However, self SCSs may still be less antigenic than nonself SCSs.

We reasoned that the relative abundance of self and nonself SCSs in a virus can be used as an indicator to evaluate the antigenicity and thus the virulence of that virus because a nonself SCS is more antigenic than a self SCS for the host immune system to avoid autoimmunity. When the number of nonself SCSs increases in a viral proteome during evolution, this means that the virus is more discoverable by the host immune system because nonself SCSs can be easily recognized as foreign objects. As a result, viral virulence may decrease. Although protein analysis methods based on SCSs have been developed since 2005 [27–33], we introduced the self/nonself concept in SCS-based analysis in a previous study for the first time [34]. In that study, we showed that when using 5 amino acid stretches as SCS units, most SCSs in the SARS-CoV-2 proteome are self SCSs in reference to the human proteome [34]. In other words, nonself SCSs are scattered in a sea of self SCSs. We also discovered nonself SCS clusters in the spike protein that may serve as an excellent candidate epitope for vaccines that efficiently induce immunity without inducing autoimmunity [34]. In the present study, we attempted to develop a simple method for predicting antigenicity based on self/nonself SCSs to understand the relationship between the Omicron variant and the human immune system.

## 2. Materials and Methods

The human reference proteome and the SARS-CoV-2 proteome sequences were obtained from NCBI (the National Center for Biotechnology Information, Bethesda, MD, USA) as described elsewhere [34]. In addition to the SARS-CoV-2 proteome reference sequence (ASM985889v3), the Delta and Omicron variant proteomes were obtained similarly (accessed on 30 November 2021 and 15 December 2021). As of 30 November 2021, there were three proteomes of the Omicron variants available at NCBI: OL698718, OL672836, and OL677199. One of them (OL698718) contained numerous X (unknown) amino acids and thus was excluded from the analysis. Because two of them (OL672836 and OL677199) had completely identical sequences, as confirmed with SIM [35] (https://web.expasy.org/sim/sim_notes.html; accessed on 27 December 2021) at Expasy (Swiss Bioinformatics Resource Portal, operated by the SIB Swiss Institute of Bioinformatics, Lausanne, Switzerland), we focused on OL672836. The Delta variant used here was OL822485, one of the most recent Delta variant proteomes available at that time.

SCSs (containing 5 amino acids) were extracted from the SARS-CoV-2 proteomes by sliding one amino acid residue at a time from the N-terminus to the C-terminus. All SARS-CoV-2 SCSs were categorized into either self (existent in the human reference proteome) or nonself (nonexistent in the human reference proteome) SCSs as described elsewhere [34]. We assigned each SCS in the SARS-CoV-2 proteome a 0 (self; invisible from the host immune system) or 1 (nonself; visible from the host immune system) at the first position of its amino acid in a protein sequence [34]. The numbers of nonself SCSs were counted and assigned in sequence maps manually based on the self (0) or nonself (1) assignments calculated by our program and exported into Microsoft Excel [34]. For analyses, ORL7b was excluded because some proteome files did not show this protein as being translated. ORF1a was also excluded because its sequence was completely redundant with that of ORF1ab.

In this study, a nonself mutation indicates a mutation that causes a self-to-nonself SCS status change. As a result of such a mutation, a nonself SCS is produced. A nonself SCS is considered antigenic by definition. Similarly, a self mutation indicates a mutation that causes a nonself-to-self SCS status change, which produces a self SCS. A self SCS is considered nonantigenic. An increase in nonself SCSs in a protein or in a proteome means a decrease in self SCSs and vice versa.

## 3. Results

We first characterized the numbers of nonself SCSs in the proteomes of the Delta and Omicron variants in comparison with that in the RefSeq (reference sequence) proteome. The numbers of nonself SCSs in the proteomes of the Delta and Omicron variants did not differ much from that in the RefSeq proteome (Table 1). The number of nonself SCSs in the Delta variant proteome increased by just one, and the number of nonself SCSs in the Omicron variant proteome decreased by just one, although these data did not indicate that there were no self/nonself status changes in these variants. The increase or decrease in the number of nonself SCSs in each protein, Δ*N*, in reference to RefSeq was not remarkable; mostly either 0 or ±1.

**Table 1.** Number of nonself SCSs in the SARS-CoV-2 reference sequence (RefSeq) and in the Delta and Omicron variants.

|  | ORF1ab | S | ORF3a | E | M | ORF6 | ORF7a | ORF8 | N | ORF10 | Total |
| --- | --- | --- | --- | --- | --- | --- | --- | --- | --- | --- | --- |
| **RefSeq (ASM985889v3)<sup>1</sup>** |  |  |  |  |  |  |  |  |  |  |  |
| Number of SCSs (*n*) | 7092 | 1269 | 271 | 71 | 218 | 57 | 117 | 117 | 415 | 34 | **9661** |
| Number of nonself SCSs (*n*) | 642 | 97 | 31 | 7 | 15 | 6 | 6 | 13 | 29 | 6 | **852** |
| Percentage (%) | 9.05 | 7.64 | 11.44 | 9.86 | 6.88 | 10.53 | 5.13 | 11.11 | 6.99 | 17.65 | **8.81** |
| **Delta (OL822485)** |  |  |  |  |  |  |  |  |  |  |  |
| Number of SCSs (*n*) | 7092 | 1267 | 271 | 71 | 218 | 57 | 117 | 115 | 415 | 34 | **9657** |
| Number of nonself SCSs (*n*) | 643 | 98 | 29 | 7 | 15 | 6 | 6 | 13 | 30 | 6 | **853** |
| Percentage (%) | 9.67 | 7.73 | 10.70 | 9.86 | 6.88 | 10.53 | 5.13 | 11.30 | 7.23 | 17.64 | **8.83** |
| $\Delta N$ (Delta–RefSeq) | **+1** | **+1** | **-2** | **0** | **0** | **0** | **0** | **0** | **+1** | **0** | **+1** |
| **Omicron (OL672836)** |  |  |  |  |  |  |  |  |  |  |  |
| Number of SCSs (*n*) | 7088 | 1266 | 271 | 71 | 218 | 57 | 117 | 117 | 412 | 34 | **9651** |
| Number of nonself SCSs (*n*) | 642 | 98 | 31 | 7 | 14 | 6 | 6 | 13 | 28 | 6 | **851** |
| Percentage (%) | 9.06 | 7.74 | 11.44 | 9.86 | 6.42 | 10.53 | 5.13 | 11.11 | 6.80 | 17.65 | **8.82** |
| $\Delta N$ (Omicron–RefSeq) | **0** | **+1** | **0** | **0** | **-1** | **0** | **0** | **0** | **-1** | **0** | **-1** |

<sup>1</sup> RefSeq data are taken from Supplementary Table S1 of Otaki et al. (2021) [34].

We next focused on the spike protein. Nonself SCSs were not concentrated in the RBD of the spike protein of the Delta variant (Figure 1). In contrast, many nonself SCSs were localized in the receptor binding domain (RBD) of the spike protein of the Omicron variant (Figure 2). Focusing on the RBD, the Delta variant had 2 nonself SCSs created by mutations (self-to-nonself status changes), and no nonself SCSs found in RefSeq disappeared (nonself-to-self status changes) (Figures 1 and 3). Thus, the net increase in nonself SCSs was +2. In contrast, the Omicron variant had 7 nonself SCSs (due to the following 7 mutations, G339D, S375F, S477N, T478K, Q498R, N501Y, and Y505H, among 15 mutations), and 3 nonself SCSs found in RefSeq disappeared (Figures 2 and 3). Thus, the net increase in nonself SCSs was +4.

**Figure 1.**
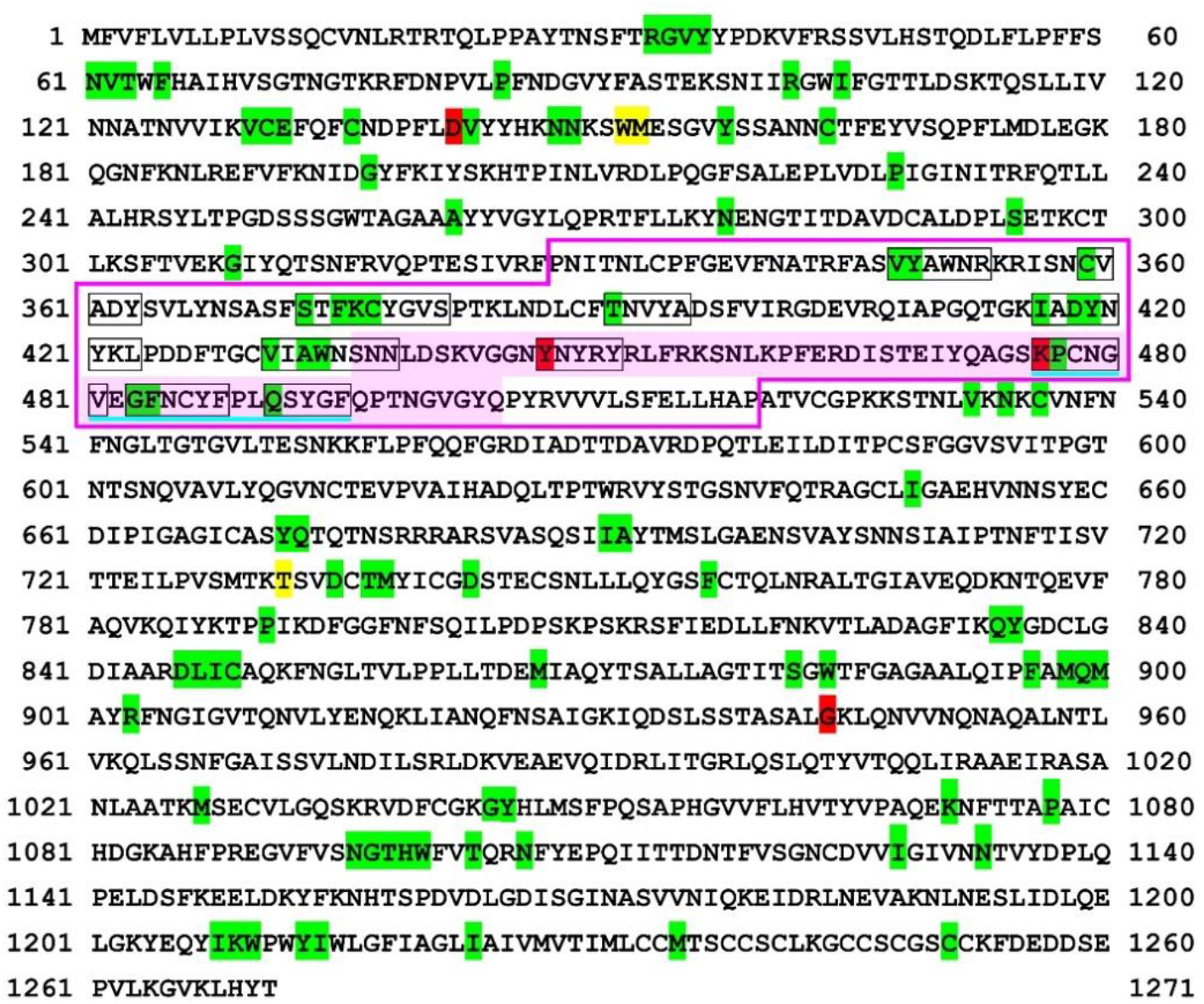
Self/nonself mapping of the SARS-CoV-2 spike protein of the Delta variant. The first amino acids of nonself SCSs are indicated by green (present in RefSeq) or red (new in this variant) shading. Other SCSs with no such indications are self SCSs. The receptor binding domain (RBD) [36] is boxed in pink lines. The receptor binding motif (RBM) [37,38] is shaded in pink. The nonself SCSs in the RBM are boxed in black lines. A potentially important nonself SCS region for vaccine development in the RBD is underlined in blue. For self/nonself mapping of RefSeq, see Otaki et al. (2021) [34].

**Figure 2.**
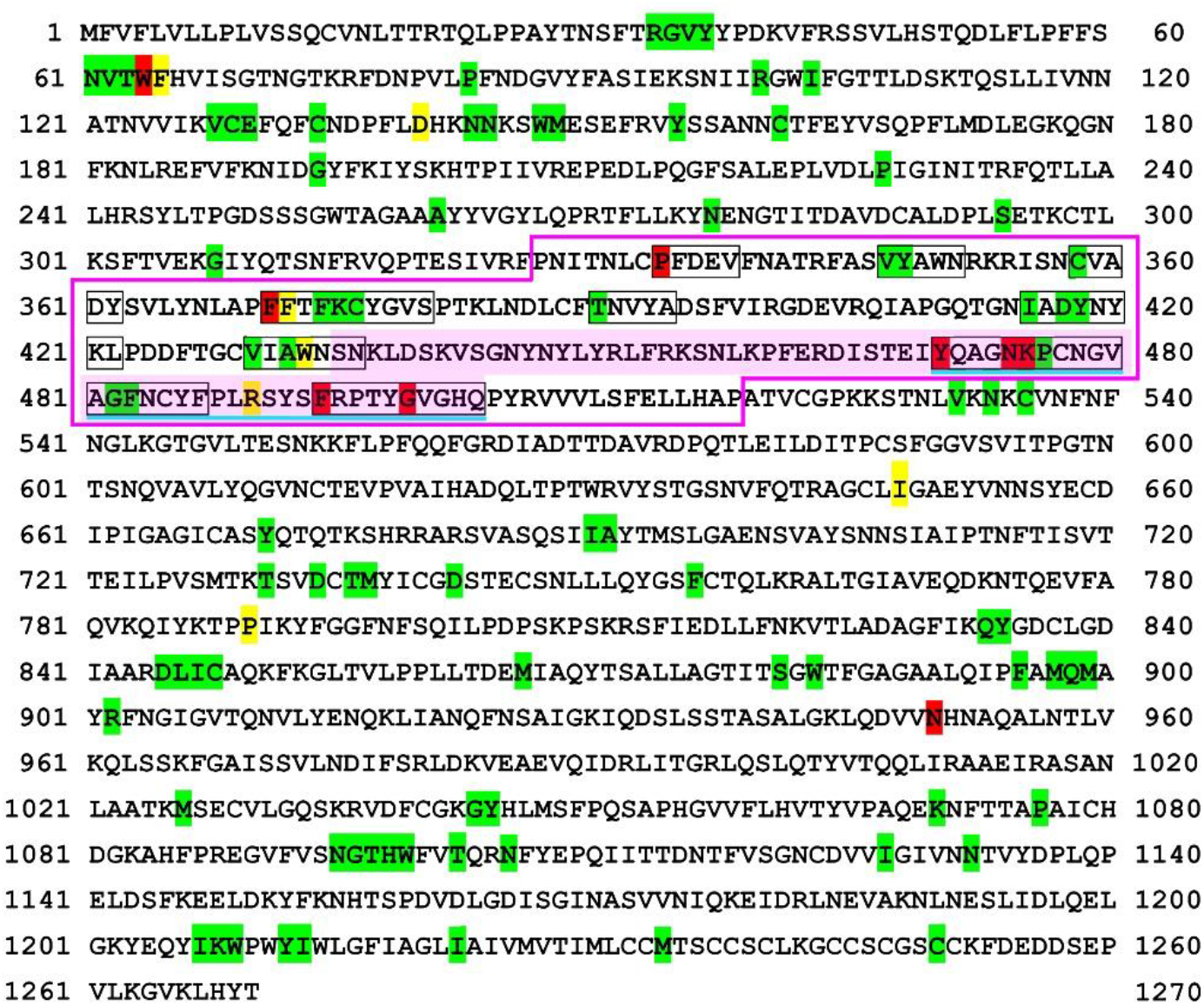
Self/nonself mapping of the SARS-CoV-2 spike protein of the Omicron variant. For colors and underlines, see legend in Figure 1. The nonself SCS at 63, TWFHV, is produced from TWFHA in RefSeq but without a status change and hence is shaded in green despite its amino acid change.

**Figure 3.**
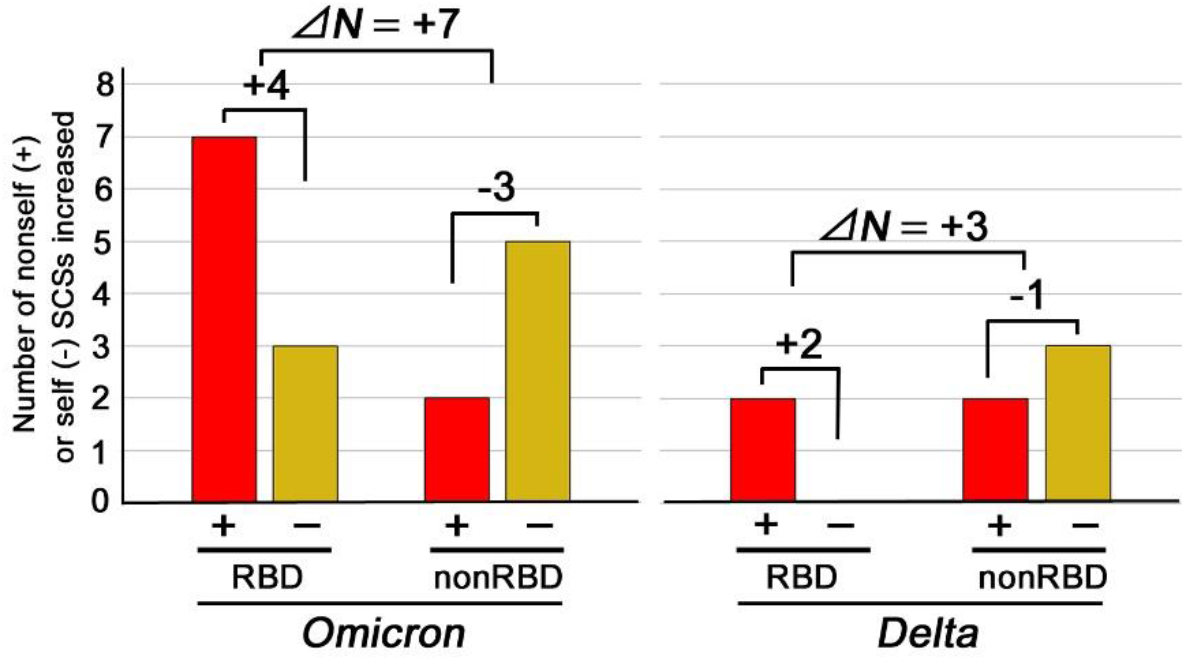
The number of nonself (+) and self (−) SCSs increased in the RBD or non-RBD regions in the Omicron and Delta variants. Newly emerged nonself SCSs (self-to-nonself status changes) are shown in red bars, whereas newly emerged self SCSs (nonself-to-self status changes) are shown in yellow bars. Note the difference between RBD and non-RBD and the difference between the Omicron and Delta variants.

Interestingly, this tendency of an increase in the number of nonself SCSs in the RBD was not observed in the non-RBD regions. Instead, the net changes in the number of nonself SCSs in the non-RBD region of the Omicron and Delta variants were −3 and −1, respectively (Figure 3). The differences in the net changes between the RBD and non-RBD regions of the Omicron and Delta variants, Δ*N*, were +7 and +3, respectively (Figure 3). In both variants, an increase in the number of nonself SCSs in the RBD was compensated for by a decrease in the number of nonself SCSs (an increase in the number of self SCSs) in the non-RBD regions (Figure 3), resulting in a net spike change of just +1 (Table 1, Figure 3).

Notably, in the receptor binding motif (RBM) of the Omicron variant, 3 novel nonself SCSs (YQAGN, NKPCN, and KPCNG) were localized immediately at the N-terminal side of the potential vaccine epitope identified in a previous study [34], extending the epitope region toward the N-terminal side (Figure 2). Furthermore, 2 novel nonself SCSs (FRPTY and GVGHQ) were localized immediately at the C-terminal side of the potential epitope, extending the epitope region toward the C-terminal side up to the end of the RBM (Figure 2). On the other hand, one nonself SCS (QSYGF) found in RefSeq and the Delta variant disappeared (Figures 1 and 2). Together, in a 34 amino acid stretch of the C-terminal end of the RBM, 8 nonself SCSs (YQAGN, NKPCN, KPCNG, PCNGV, GFNCY, FNCYF, FRPTY, and GVGHQ) were concentrated tandemly.

## 4. Discussion

In the present study, we reasoned that the number of nonself SCSs in a proteome or in a protein is a determinant of its antigenicity. We further reasoned that antigenicity is probably reflected directly in the virulence of the virus. Not only the number but also the locations of nonself SCSs in a protein (e.g., in a binding domain, at an active site, or in proximity to one another) would matter to determine the antigenicity of nonself SCSs.

Based on the above logic, we endeavored to urgently characterize the Omicron variant of SARS-CoV-2. Although immunological validity should be evaluated in further studies, this study proposes a novel method of predicting viral virulence based on the amino acid sequences of viral proteins.

Theoretically, a virus evolves under selection pressure for higher infectivity and transmissibility, and in this evolutionary process, the amino acid residues of the binding site for its receptor are mutated for higher affinity. This type of evolution is here called “offensive” evolution. Offensive evolution may attenuate when any functional disadvantage unavoidably associated with excessive mutations for higher affinity overwhelms the mutational functional advantage. Offensive evolution may also be attenuated when nonself SCSs accumulate for higher antigenicity.

Alternatively, a virus theoretically evolves for higher sequence mimicry to the host SCS repertoire defined by the host proteome [34], and in this evolutionary process, nonself SCSs decrease in number for lower antigenicity. This type of evolution is here called “defensive” evolution. Defensive mimicry of host SCSs by increasing self SCSs is advantageous for viruses because they cannot be readily detected by the host immune system. Excessive mimicry evolution may also be attenuated due to an unavoidable functional compromise.

Given a sufficient period of time, an equilibrium state between offensive and defensive evolution may be established through a functional compromise of a virus. However, in reality, a virus may evolve within a limited period (i.e., replications). Either offensive or defensive evolution may settle at local maxima of survival, depending on various environmental and host conditions.

We discovered that the Omicron variant has accumulated self-to-nonself status change mutations at the RBD of the spike protein. The Omicron mutant had 7 new nonself SCSs in the RBD, in contrast to just 2 additions in the Delta variant. The net increase in nonself SCSs within the RBD was +4 in the Omicron variant, in contrast to +2 in the Delta variant. This increase is likely immunologically significant, considering that the RBD is readily accessible to other proteins, such as ACE2, antibodies and T-cell receptors. Interestingly, there were just 2 newly added nonself SCSs in the non-RBD regions in both the Omicron and Delta variants, but in the Omicron variant, more new self SCSs were introduced in the non-RBD regions compared with the non-RBD regions in the Delta variant and the RBD of the Omicron variant. This increase in self SCSs in the non-RBD regions in the Omicron variant may compensate for the increase in nonself SCSs in the RBD to make the net nonself SCS increase small for lower antigenicity. Indeed, the differences in nonself SCS changes between the RBD and non-RBD regions, Δ*N*, were +7 and +3 in the Omicron and Delta variants, respectively.

It seems that for the Omicron variant, the selection pressure for higher binding affinity to ACE2 (i.e., higher infectivity and transmissibility) is so large that an increase in the number of nonself SCSs in the RBD was unavoidably allowed despite its immunological disadvantage for the virus. In other words, an increase in nonself SCSs in the RBD likely means a compromise of the virus, which probably indicates low virulence. In support of this view, nonself mutations in the RBD appear to contribute directly to higher affinity to ACE2 [18]. Therefore, the Omicron variant is probably a product of offensive evolution but with a compromise of high antigenicity. The Delta variant may also be a product of offensive evolution but to a lesser degree. Due to relatively high antigenicity, the current offensive evolution of the Omicron variant may attenuate at some point.

Based on the idea that SCS frequencies in proteomes are related to the phylogenetic and parasitic status of organisms [39,40], we speculated previously that the SARS-CoV-2 proteome will eventually evolve to accumulate self SCSs to defray molecular and cellular attack from the human immune system [34]. However, this defensive evolution based on the sequence mimicry hypothesis was not detected clearly in the Omicron and Delta variants. An increase in self SCS was detected only in the non-RBD regions in the Omicron and Delta variants. Gears may be changed at some point toward a defensive mode of evolution, but offensive evolution may settle at local maxima of survival for long periods of time.

In addition to the above discussion on the number of nonself SCSs, further discussion can be made in terms of the location of nonself SCSs within the RBM in the RBD. We discovered a candidate stretch of amino acids that contained nonself SCSs within the RBM of the RefSeq spike protein, a potential epitope for vaccine development to avoid vaccine-induced autoimmunity [34]. Interestingly, in the Omicron variant but not in the Delta variant, 3 additional nonself SCSs (YQAGN, NKPCN, and KPCNG) are present at the N-terminal side of the potential epitope region, and furthermore, 2 additional nonself SCSs (FRPTY and GVGHQ) are present at the C-terminal side, although one nonself SCS in RefSeq is not present. Together, the candidate epitope region in RefSeq has been extended to both the N-terminal and C-terminal sides. Because this region in the RBM is important for ACE2 binding, the expansion of this nonself SCS epitope region here was probably unavoidable to increase the binding affinity for ACE2 at the expense of an increase in antigenicity. This result supports the previous finding that this region is likely a good (probably the best) epitope candidate for vaccine development [34].

At first glance, the present results may not seem to be consistent with recent studies on the Omicron variant, which suggested an increase in virulence [10–22]. These studies evaluated whether the pre-existing immunological memory induced by previous vaccines or infection continues to be effective against the Omicron variant, but in the present study, the antigenicity of the Omicron variant itself was evaluated. The present study suggests that the Omicron variant has reduced virulence because of its relatively high antigenicity. Therefore, the Omicron variant may not cause severe symptoms. However, due to its high infectivity and transmissibility, as suggested by other studies [10–22], the Omicron variant should still be regarded with alarm.

Antigenicity defined by nonself SCSs is just a single factor to explain virulence. A virus with low virulence might preferentially affect tissues/organs such as the digestive tract and testes, leading to non-life-threatening but long-term effects, rather than tissues/organs such as the lungs, the effects of which could be life-threatening. For example, infection may cause infertility if testicular cells expressing ACE2 are preferentially infected. Moreover, due to high infectivity and transmissibility, even if virulence is low, the absolute number of hospitalized people may not decrease in the Omicron pandemic in comparison with the Delta pandemic. Therefore, the Omicron variant should be considered highly threatening in terms of public health.

The frequency of random mutations is directly proportional to the number of replications (i.e., the number of infected people), but selection pressure shapes the direction of viral evolution. The higher binding affinity to ACE2 and higher antigenicity together caused by the accumulated mutations in the Omicron variant suggest that a hampered transmission state despite a large number of replications was a driving force for the evolution of the Omicron variant. Ironically, such an unnatural state was created by worldwide vaccination, which might have helped the emergence of the Omicron variant, as suggested by a recent study [20].

## 5. Conclusions

Through an application of the SCS concept to the human-SARS-CoV-2 system together with immunological self/nonself considerations, in the present study, a novel method was used to characterize the Omicron and Delta variants. It appears that the Omicron variant increased its infectivity and transmissibility at the expense of higher antigenicity and lower virulence. Thus, the symptoms of individuals infected with the Omicron variant may be less severe than those of individuals infected with the Delta variant. We also confirmed that a specific stretch in the RBM is a good candidate epitope for vaccines. Since SARS-CoV-2 evolution might have been driven by selection pressure imposed by worldwide vaccination, vaccination-focused strategies against the Omicron variant may further enhance a current mode of evolution. Alternatives to vaccination-focused strategies [41–47] may also be useful. Continuous caution regarding the Omicron variant is necessary.

## Author Contributions

Conceptualization, J.M.O.; methodology, W.N. and M.N.; software, W.N. and M.N.; validation, J.M.O.; formal analysis, J.M.O. and W.N.; investigation, J.M.O., W.N. and M.N.; resources, W.N. and M.N.; data curation, J.M.O., W.N. and M.N.; writing—original draft preparation, J.M.O.; writing—review and editing, J.M.O.; visualization, J.M.O.; supervision, J.M.O. and M.N.; project administration, J.M.O.; funding acquisition, J.M.O. and M.N. All authors have read and agreed to the published version of the manuscript.

## Institutional Review Board Statement

Not applicable.

## Informed Consent Statement

Not applicable.

## Data Availability Statement

All data that support the conclusions of the study are included in this paper and the related Supplementary Information. The source codes for human SCS analysis and for SARS-CoV-2 SCS analysis are freely available at https://adslab-uryukyu.github.io/scs-sars-cov-2/.

## Acknowledgments

The authors would like to acknowledge Mr. Tatsuya Ishibashi for confirming self/nonself assignments using an independent program of his own and Prof. Hideo Yamasaki, Dr. Wataru Taira, and other laboratory members of the BCPH Unit of Molecular Physiology for discussion.

## References

1. Zhou, P. et al. A pneumonia outbreak associated with a new coronavirus of probable bat origin. Nature 2020, 579, 270–273. DOI: 10.1038/s41586-020-2012-7.

2. Wu, F. et al. A new coronavirus associated with human respiratory disease in China. Nature 2020, 579, 265–269. DOI: 10.1038/s41586-020-2008-3.

3. Zhang, X. et al. Viral and host factors related to the clinical outcome of COVID-19. Nature 2020, 583, 437–440. DOI: 10.1038/s41586-020-2355-0.

4. Zhang, Z. et al. The establishment of reference sequence for SARS-CoV-2 and variation analysis. J. Med. Virol. 2020, 92, 667–674. DOI: 10.1002/jmv.25762.

5. South African Institute for Communicable Diseases (NICD) Division of the National Health Laboratory Service. New COVID-19 variant detected in South Africa. https://www.nicd.ac.za/new-covid-19-variant-detected-in-south-africa/ Accessed on 20 December 2021.

6. World Health Organization. Tracking SARS-CoV-2 variants. https://www.who.int/en/activities/tracking-SARS-CoV-2-vari-ants/ Updated on 13 December 2021. Accessed on 20 December 2021.

7. European Centre for Disease Prevention and Control (ECDC). Threat assessment brief: Implications of the further emergence and spread of the SARS CoV 2 B.1.1.529 variant of concerm (Omicron) for the EU/EEA first update. https://www.ecdc.europa.eu/en/publications-data/covid-19-threat-assessment-spread-omicron-first-update Accessed on 20 December 2021.

8. Espenhain, L. et al. Epidemiological characterization of the first 785 SARS-CoV-2 Omicron variant cases in Denmark, December 2021. Euro Surveill. 2021 26. DOI: 10.2807/1560-7917.ES.2021.26.50.2101146.

9. CDC COVID-19 Response Team. SARS-CoV-2 B.1.1.529 (Omicron) Variant—United States, December 1–8, 2021. MMWR Morb. Mortal. Wkly Rep. 2021, 70, 1731–1734. DOI: 10.15585/mmwr.mm7050e1.

10. Ingraham, N.E.; Ingbar, D.H. The omicron variant of SARS-CoV-2: Understanding the known and living with unknown. Clin. Transl. Med. 2021, 11, e685. DOI: 10.1002/ctm2.685.

11. Kandeel, M.; Mohamed, M.E.M.; Abd El-Lateef, H.M.; Venugopala, K.N.; El-Beltagi, H.S. Omicron variant genome evolution and phylogenetics. J. Med. Virol. 2021. Online ahead of print. DOI: 10.1002/jmv.27515.

12. Wang, L.; Cheng, G Sequence analysis of the Emerging Sars-CoV-2 Variant Omicron in South Africa. J. Med. Virol. 2021. Online ahead of print. DOI: 10.1002/jmv.27516.

13. Kannan, S.R.; Spratt, A.N.; Sharma, K.; Chand, H.S.; Byrareddy, S.N.; Singh, K. Omicron SARS-CoV-2 variant: Unique features and their impact on pre-existing antibodies. J. Autoimmun. 2021, 126, 102779. DOI: 10.1016/j.jaut.2021.102779.

14. Saxena, S.; Kumar, S.; Ansari, S.; Paweska, J.T.; Maurya, V.K.; Tripathi, A.K.; Abdel-Moneim, A. Characterization of the novel SARS-CoV-2 Omicron (B.1.1.529) Variant of Concern and its global perspective. J. Med. Virol. 2021. Online ahead of print. DOI: 10.1002/jmv.27524.

15. Kumar, S.; Thambiraja, T.S.; Karuppanan, K.; Subramaniam, G. Omicron and Delta variant of SARS-CoV-2: A comparative computational study of spike protein. J. Med. Virol. 2021. Online ahead of print. DOI: 10.1002/jmv.27526.

16. Pawłowski, P. SARS-CoV-2 variant Omicron (B.1.1.529) is in a rising trend of mutations increasing the positive electric charge in crucial regions of the spike protein S. Acta Biochim. Pol. 2021. Online ahead of print. DOI: 10.18388/abp.2020_6072.

17. Pascarella, S. Ciccozzi, M.; Bianchi, M.; Benvenuto, D.; Cauda, R.; Cassone, A. The electrostatic potential of the Omicron variant spike is higher than in Delta and Delta-plus variants: a hint to higher transmissibility? J. Med. Virol. 2021. Online ahead of pint. DOI: 10.1002/jmv.27528.

18. Chen, J.; Wang, R.; Gilby, N.B.; Wei, G.-W. Omicron (B.1.1.529): Infectivity, vaccine breakthrough, and antibody resistance. ArXiv 2021. arXiv:2112.01318v1.

19. Miller, N.L.; Clark, T.; Raman, R.; Sasisekharan, R. Insights on the mutational landscape of the SARS-CoV-2 Omicron variant. bioRxiv 2021. 2021.12.06.471499. DOI: 10.1101/2021.12.06.471499.

20. Wang, R.; Chen, J.; Wei, G.-W. Mechanisms of SARS-CoV-2 evolution revealing vaccine-resistant mutations in Europe and America. J. Phys. Chem. Lett. 2021, 12, 11850–11857. DOI: 10.1021/acs.jpclett.1c03380.

21. Zhang, L. et al. The significant immune escape of pseudotyped SARS-CoV-2 variant Omicron. Emerg. Microbes Infect. 2021. Online ahead of print. DOI: 10.1080/22221751.2021.2017757.

22. Cele, S. et al. SARS-CoV-2 Omicron has extensive but incomplete escape of Pfizer BNT162b2 elicited neutralization and requires ACE2 for infection. medRxiv 2021. 2021.12.08.21267417. DOI: 10.1101/2021.12.08.21267417.

23. Redd, A.D. et al. Minimal cross-over between mutations associated with Omicron variant of SARS-CoV-2 and CD8^+^ T cell epitopes identified in COVID-19 convalescent individuals. bioRxiv 2021. 2021.12.06.471446.

24. Bjorkman, P. J.; Saper, M. A.; Samraoui, B.; Bennett, W. S.; Strominger, J. L.; Wiley, D. C. Structure of the human class I histocompatibility antigen, HLA-A2. Nature 1987, 329, 506–512. DOI: 10.1038/329506a0.

25. Rossjohn, J.; Gras, S.; Miles, J. J.; Turner, S. J.; Godfrey, D. I.; McCluskey, J. T cell antigen receptor recognition of antigen-presenting molecules. Annu. Rev. Immunol. 2015, 33, 169–200. DOI: 10.1146/annurev-immunol-032414-112334.

26. Theodossis, A. et al. Constraints within major histocompatibility complex class I restricted peptides: Presentation and consequences for T-cell recognition. Proc. Natl. Acad. Sci. USA 2010, 107, 5534–5539. DOI: 10.1073/pnas.1000032107.

27. Otaki, J. M.; Ienaka, S.; Gotoh, T.; Yamamoto, H. Availability of short amino acid sequences in proteins. Protein Sci. 2005, 14, 617–625. DOI: 10.1110/ps.041092605.

28. Otaki, J. M.; Gotoh, T.; Yamamoto, H. Potential implications of availability of short amino acid sequences in proteins: an old and new approach to protein decoding and design. Biotechnol. Annu. Rev. 2008, 14, 109–141. DOI: 10.1016/S1387-2656(08)00004-5.

29. Otaki, J. M.; Tsutsumi, M.; Gotoh, T.; Yamamoto, H. Secondary structure characterization based on amino acid composition and availability in proteins. J. Chem. Inf. Model. 2010, 50, 690–700. DOI: 10.1021/ci900452z.

30. Tsutsumi, M.; Otaki, J. M. Parallel and antiparallel β-strands differ in amino acid composition and availability of short constituent sequences. J. Chem. Inf. Model. 2011, 51, 1457–1464. DOI: 10.1021/ci200027d.

31. Motomura, K.; Fujita, T.; Tsutsumi, M.; Kikuzato, S.; Nakamura, M.; Otaki, J. M. Word decoding of protein amino acid sequences with availability analysis: a linguistic approach. PLoS One 2012, 7: e50039. DOI: 10.1371/journal.pone.0050039.

32. Motomura, K.; Nakamura, M.; Otaki, J.M. A frequency-based linguistic approach to protein decoding and design: Simple concepts, diverse applications, and the SCS Package. Comput. Struct. Biotechnol. J. 2013, 5, e201302010. DOI: 10.5936/csbj.201302010.

33. Endo, S.; Motomura, K.; Tsuhako, M.; Kakazu, Y.; Nakamura, M., Otaki, J. M. Search for human-specific proteins based on availability scores of short constituent sequences: Identification of a WRWSH protein in human testis. In Computational Biology and Chemistry; Behzadi, P., Bernabò, N., Eds.; IntechOpen: London, UK, 2019; pp. 11–33. DOI: 10.5772/intechopen.89653.

34. Otaki, J.M.; Nakasone, W.; Nakamura, M. Self and nonself short constituent sequences of amino acids in the SARS-CoV-2 proteome for vaccine development. COVID 2021, 1, 555–574. DOI: 10.3390/covid1030047.

35. Huang, X.; Miller, W. A time-efficient, linear-space local similarity algorithm. Adv. Appl. Math. 1991, 12, 337–357.

36. Zhang B. et al. Mining of epitopes on spike protein of SARS-CoV-2 from COVID-19 patients. Cell Res. 2020, 30, 702–704. DOI: 10.1038/s41422-020-0366-x.

37. Lan, J. et al. Structure of the SARS-CoV-2 spike receptor-binding domain bound to the ACE2 receptor. Nature 2020, 581, 215–220. DOI: 10.1038/s41586-020-2180-5.

38. Wrapp, D.; Wang, N.; Corbett, K. S.; Goldsmith, J. A.; Hsiech, C.-L.; Abiona, O.; Graham, B. S.; McLellan, J. S. Cryo-EM structure of the 2019-nCoV spike in the prefusion conformation. Science 2020, 367, 1260–1263. DOI: 10.1126/science.abb2507.

39. Pe’er, I.; Felder, C. E.; Man, O.; Silman, I.; Sussman, J. L.; Beckmann, J. S. Proteomic signatures: Amino acid and oligopeptide compositions differentiates among phyla. Proteins 2004, 54, 20–40. DOI: 10.1002/prot.10559.

40. Zemková, M.; Zahradní, D.; Mokrejš, M.; Flegr, J. Parasitism as the main factor shaping peptide vocabularies in current organisms. Parasitology 2017, 144, 975–983. DOI: 10.1017/S0031182017000191.

41. Singh, T.U.; Parida, S.; Lingaraju, M.C.; Kesavan, M.; Kumar, D.; Singh, R.K. Drug repurposing approach to fight COVID-19. Pharmacol. Rep. 2020, 72, 1479–1508. DOI: 10.1007/s43440-020-00155-6.

42. Chugh, H.; Awasthi, A.; Agarwal, Y.; Gaur, R.K.; Dhawan, G.; Chandra, R. A comprehensive review on potential therapeutics interventions for COVID-19. Eur. J. Pharmacol. 2021, 890, 173741. DOI: 10.1016/j.ejphar.2020.173741.

43. Majumder, J.; Minko, T. Recent development on therapeutic and diagnostic approaches for COVID-19. AAPS J. 2021, 23, 14. DOI: 10.1208/s12248-020-00532-2.

44. Gavriatopoulou, M. et al. Emerging treatment strategies for COVID-19 infection. Clin. Exp. Med. 2021, 21, 167–179. DOI: 10.1007/s10238-020-00671-y.

45. Yamasaki, H. Blood nitrate and nitrite modulating nitric oxide bioavailability: Potential therapeutic functions in COVID-19. Nitric Oxide 2020, 103, 29–30. DOI: 10.1016/j.niox.2020.07.005.

46. Yang, Y.; Islam, M. S.; Wang, J.; Li, Y.; Chen, X. Traditional Chinese Medicine in the treatment of patients infected with 2019-new coronavirus (SARS-CoV-2): A review and perspective. Int. J. Biol. Sci. 2020, 16, 1708–1717. DOI: 10.7150/ijbs.45538.

47. Liu, M.; Gao, Y.; Yuan, Y.; Yang, K.; Shi, S.; Zhang, J.; Tian, J. Efficacy and safety of Integrated Traditional Chinese and Western Medicine for corona virus disease 2019 (COVID-19): a systematic review and meta-analysis. Pharmacol. Res. 2020, 158, 104896. DOI: 10.1016/j.phrs.2020.104896.

